# cGAS activation during human cytomegalovirus infection is driven by exogenous DNA

**DOI:** 10.64898/2026.03.27.714697

**Authors:** Matin Mahmoudi, Yao-Tang Lin, Michael Nevels, Finn Grey

## Abstract

Type I interferon (IFN) induction is a central component of the innate immune response to viral infection, and the cytosolic DNA sensor cyclic GMP-AMP synthase (cGAS) has been identified as a key mediator of IFN production during human cytomegalovirus (HCMV) infection. However, how cGAS detects HCMV remains unresolved, as the viral genome is encapsidated and trafficked directly to the nucleus, limiting cytoplasmic exposure. Here, we show that IFN induction during HCMV infection of primary fibroblast cells is predominantly driven by cGAS recognition of exogenous DNA present in standard laboratory virus preparations rather than the encapsidated viral genome. DNase treatment of AD169 and low-passage TB40/E-GFP viral stocks substantially reduced total DNA content without affecting infectivity, yet markedly abrogated IFN induction, IFN-stimulated gene expression and IRF3 nuclear translocation. Immunofluorescence analysis further revealed cytoplasmic accumulation of DNA in cells infected with untreated virus stocks, which was absent following DNase treatment. Together, these findings demonstrate that contaminating DNA in viral preparations is sufficient to activate cGAS and drive IFN responses during HCMV infection *in vitro*, highlighting a critical confounding factor in studies of innate immune sensing.

**Author Summary:** Human cytomegalovirus (HCMV) is a common herpesvirus that establishes lifelong infection and can cause serious disease in immunocompromised individuals and newborns. When cells detect viral infection, they produce type I interferons (IFNs), antiviral molecules that help limit virus spread. Previous studies have suggested that HCMV is sensed by a cellular DNA sensor called cGAS, which detects viral DNA in the cytoplasm and triggers IFN production. However, how cGAS gains access to the HCMV genome has remained unclear, because the viral DNA is enclosed within a protective capsid and transported directly to the nucleus during infection.

In this study, we show that most IFN production observed during HCMV infection of fibroblast cells *in vitro* is driven not by sensing of the viral genome itself, but by contaminating DNA present in standard laboratory virus preparations. Treating virus stocks with DNase to remove this exogenous DNA abolished IFN induction without affecting viral infectivity. These findings highlight the importance of controlling for exogenous nucleic acids when interpreting how host cells detect viral infection.

## Introduction

Cytokines are a critical first line of defence against infections by shaping the innate and adaptive immune responses [1, 2]. Production is triggered through recognition of pathogen-associated molecular patterns (PAMPs) by pattern recognition receptors (PRRs), including Toll-like receptors, RIG-I-like receptors, and cytosolic DNA sensors [3]. PAMPs include structures such as lipopolysaccharides, double-stranded RNA, and cytoplasmic DNA. Detection triggers signalling cascades that drive the translocation of transcription factors like IRF3 and NF-κB into the nucleus, inducing cytokine and chemokine expression. Cytokines, including interferons (IFN), subsequently activate their receptors, upregulating hundreds of genes (e.g., IFN-stimulated genes (ISGs)) to establish an antiviral state [4].

*In vitro* infections with the β-herpesvirus HCMV elicits a strong IFN response [5-9]. The cytosolic DNA sensor cyclic GMP-AMP synthase (cGAS) has been identified as the primary sensor of HCMV, as its loss or inhibition abrogates IFN production in several cell types [10, 11]. cGAS recognises double-stranded DNA (dsDNA) and synthesises 2′-3′-cyclic GMP-AMP (cGAMP) from ATP and GTP in response [12]. cGAMP then activates the adaptor protein STING that subsequently translocates from the ER to the Golgi, where it recruits TBK1 [13, 14]. Here, TBK1 activates IRF3 by phosphorylation [15, 16]. The mechanism by which cGAS detects HCMV remains unresolved, as its DNA is encapsidated and transported directly to the nuclear pore complex, seemingly avoiding cytoplasmic exposure. Possible explanations for cytoplasmic detection of HCMV include release of viral DNA from defective particles and capsid destabilization by host proteins, analogous to the role of TRIM5α during HIV-1 infection [17-20]. MxB has been suggested to play a role in herpesvirus capsid disruption [21].

An alternative explanation is stimulation of cGAS by contaminating DNA produced during viral preparation triggers IFN production. Here, we show that exogenous DNA present in standard laboratory virus preparations is the predominant trigger of cGAS activation during HCMV infection of fibroblast cells. These findings call for a re-evaluation of how IFN induction during HCMV infection is interpreted in *in vitro* systems.

## Results

Previous reports identified cGAS as the primary PRR responsible for triggering IFN production following HCMV infection [10, 11]. To confirm these results, siRNA-mediated knockdown of cGAS was performed in human primary fibroblast cells and confirmed by RT-qPCR (Figure 1A). Consistent with previous results, IFN production in response to HCMV was completely abrogated in knockdown cells (Figure 1B). cGAS knockdown also resulted in almost complete loss of IFNB1 mRNA (Figure 1C) and reduced IRF3 nuclear translocation (Figure 1D), supporting previous publications identifying cGAS as responsible for HCMV detection in fibroblast cells [10, 11].

**Figure 1.**
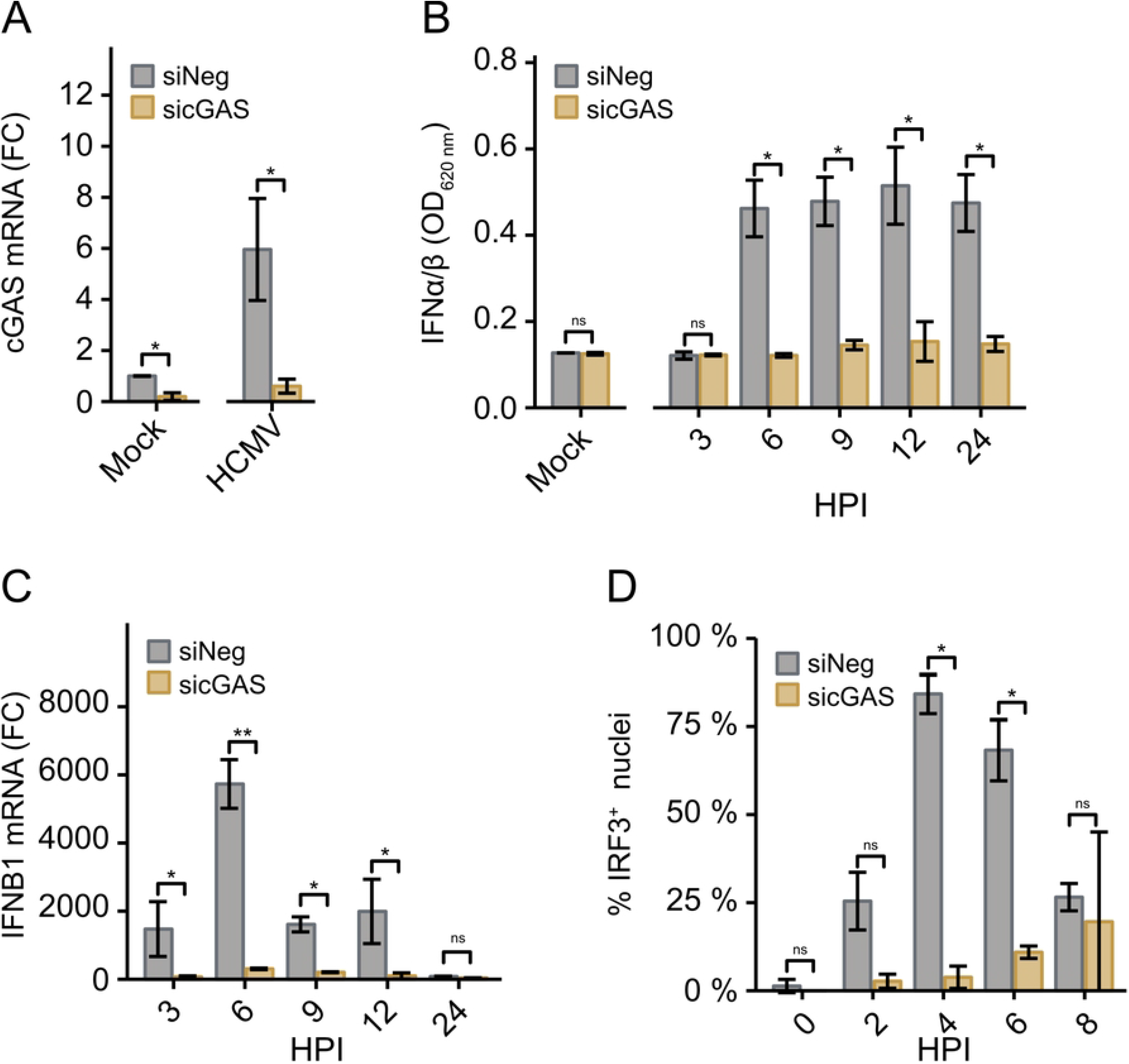
HCMV induces cGAS-mediated immune response in HDFn cells. Human dermal fibroblast cells were transfected with cGAS-targeting siRNA (sicGAS) and infected with HCMV strain TB40/E-GFP at an MOI of 3, 48 h post transfection. (A) RNA was harvested 9 hpi and cGAS mRNA levels quantified by RT-qPCR and normalised to mock-infected control cells. (B) Supernatants were assessed for secreted type I IFNs using the HEK-Blue IFN-α/β cell line. OD_620_ refers to absorbance following incubation of HEK-Blue supernatants with QUANTI-Blue reagent. (C) RNA was harvested at the indicated time points and IFNB1 mRNA levels quantified by RT-qPCR and normalised to mock-infected control cells. (D) Cells were fixed at the indicated time points using 4% paraformaldehyde and permeabilised with 0.1 % Triton X-100. Cells were then probed with anti-IRF3 antibody (Cell Signaling Technology, clone D6I4C) prior to staining with anti-Rabbit Alexa Fluor 568 (Invitrogen, A10037). An average of 30 cells were counted per condition for each independent experiment. *Error bars: standard deviation. Students t test (A) or Two*-*way ANOVA (B, C and D) was performed on all data (n = 3): ns, not significant; * p < 0*.*05; ** p < 0*.*005; *** p < 0*.*0005*.

However, as the HCMV genome is protected during transit to the nuclear core complex, it remains unclear how cGAS is triggered. We hypothesised that recognition of exogenous DNA in viral preparations may stimulate cGAS. To test this, viral preparations of lab-adapted (AD169) and low-passage (TB40/E-GFP) HCMV strains, were treated with DNase before infection of fibroblast cells. Viral stocks were generated using a standard protocol (see Materials and Methods) and split into two fractions, one of which was DNase treated (Figure 2A). DNase treatment should selectively degrade non-encapsidated DNA, while encapsidated viral genomes remain protected by the capsid and envelope.

**Figure 2.**
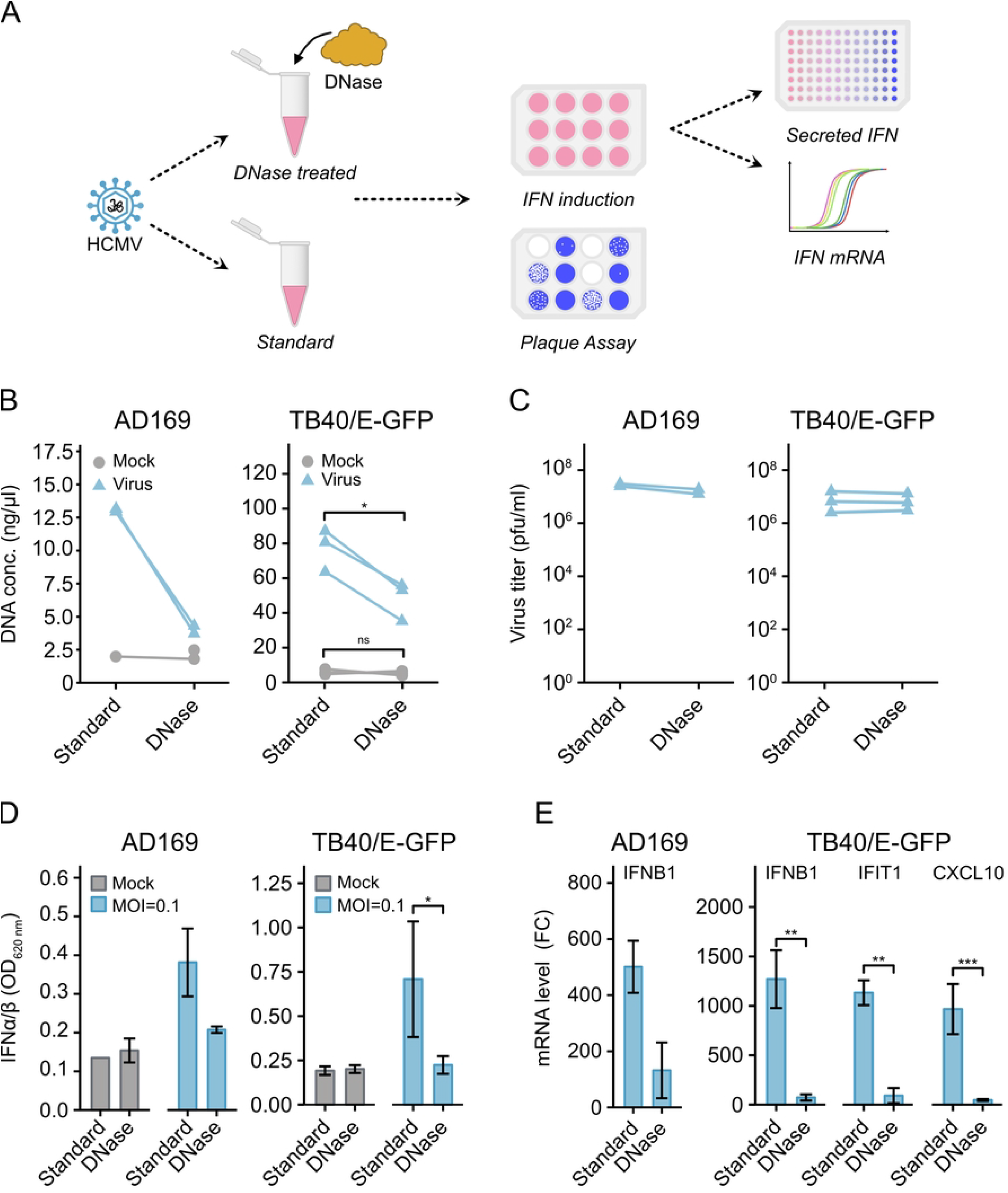
DNase treatment reduces immunogenicity of HCMV stocks. Stocks of HCMV strain AD169 and TB40E-GFP were treated with TURBO DNase and used to infect human dermal fibroblast cells. (A) Schematic of experimental setup. (B) Virus stocks were assessed for DNA concentration by Qubit assay and (C) infectivity by plaque assay with or without DNase treatment. (D) Fibroblast cells were infected at an MOI of 0.1 and supernatants were harvested at 6 hpi and assessed for secreted type I IFNs using the HEK-Blue IFN-α/β cell line. OD_620 nm_ refers to absorbance following incubation of HEK-Blue supernatants with QUANTI-Blue reagent. (E) RNA was harvested at 6 HPI and analysed by RT-qPCR to determine IFNB1, IFIT1 and CXCL10 mRNA levels. *Error bars: standard deviation. Student’s t test (B and C) or One way ANOVA (D) was performed on data from TB40/E-GFP infections (n=3). AD169 (n=2): ns, not significant; * p < 0*.*05; ** p < 0*.*005; *** p < 0*.*0005*.

DNase treatment significantly reduced total DNA levels in viral stocks as measured by Qubit analysis, suggesting a reduction in exogenous DNA (Figure 2B). To test whether DNase treatments impacted infectivity, the viral stocks were subjected to plaque assay analysis. Titres of DNase treated and untreated stocks were equivalent, indicating that treatment had little impact on infectivity (Figure 2C).

To test the effect of DNase treatment on IFN production, fibroblast cells were infected with treated or untreated virus or mock at an MOI of 0.1. While infection with untreated virus resulted in robust IFN production, DNase treatment of viral stocks reduced IFN production to basal levels (Figure 2D). Gene expression analysis using RT-qPCR showed that DNase treatment reduced IFN and ISG mRNA expression close to basal levels (Figure 2E). These data indicate that IFN production is predominantly driven by recognition of exogenous DNA.

To understand the impact of exogenous DNA on IFN induction in individual cells, IRF3 activation and viral IE protein expression was examined by immunofluorescence microscopy. Infections with untreated virus stocks resulted in rapid translocation of IRF3 to the nucleus in the majority of cells, whereas DNase treatment resulted in few cells with nuclear IRF3 (Figure 3A). Specifically, infections with untreated virus resulted in nuclear IRF3 in both IE^-^ and IE^+^ cells (Figure 3B), whereas nuclear IRF3 was rare in IE^-^ cells inoculated with DNase treated virus. Furthermore, inoculation with DNase treated virus resulted in the majority of IE^+^ cells being absent of nuclear IRF3, suggesting that exogenous DNA induces IFN production in both infected and uninfected cells (Figure 3C).

**Figure 3.**
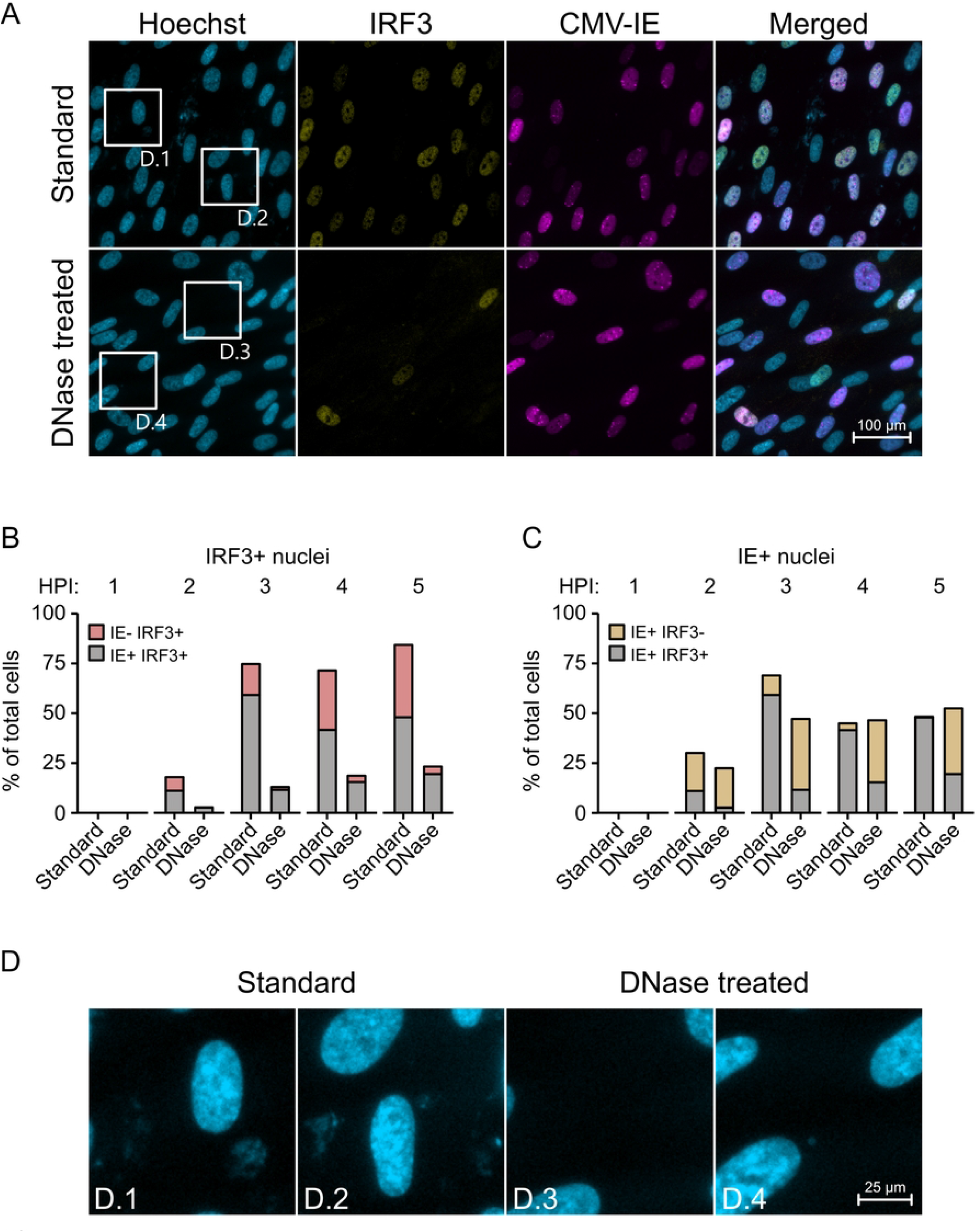
Non-infectious DNA induces IRF3 translocation during HCMV infection. Stocks of HCMV strain AD169 were treated with TURBO DNase and used to infect human dermal fibroblast cells at an MOI of 3. Cells were fixed using 4 % paraformaldehyde and permeabilised with 0.1 % Triton X-100. Cells were then probed with anti-IRF3 antibody (Cell Signaling Technology, clone D6I4C) and anti CMV-IE antibody (Millipore, clone 6F8.2), prior to staining with anti-Rabbit Alexa Fluor 488 (Invitrogen, A21206) and anti-Mouse Alexa Fluor 647 (Invitrogen, A21235). (A) Fluorescent images of cells fixed at 4 HPI. Regions within the squares are magnified in panel D. (B and C) Quantification of IE and IRF3 staining from multiple time points post infection. For quantification, all cells within an image were counted. The number of cells ranged from 179 to 349 per image. (D) Magnified panels showing presence or absence of DNA staining in the cytoplasm of cells infected with untreated virus stocks (D.1 and D.2) or DNase treated virus stocks (D.3 and D.4). Chi-squared *was performed on data, based on categorisation into four groups (IE*^*-*^*IRF3*^*-*^, *IE*^*+*^*IRF3*^*-*^, *IE*^*-*^*IRF3*^*+*^ *and IE*^*+*^*IRF3*^*+*^: ^*ns*^ *not significant, * p < 0*.*05, ** p < 0*.*005, *** p < 0*.*0005*.

Strikingly, examination of Hoechst staining revealed non-nuclear DNA in cells infected with untreated virus stocks (Figure 3D.1 and 3D.2). No such signal was apparent in infections with DNase treated stocks (Figure 3D.3 and 3D.4), reaffirming the existence of exogenous DNA during virus infection.

## Discussion

Type I IFN induction is widely interpreted as evidence of innate immune sensing during HCMV infection, with the cGAS–STING pathway proposed as the primary mediator of this response [10, 11]. However, given encapsidation of the viral genome during cytoplasmic translocation [22], the mechanism behind cGAS-detection of HCMV in the cytoplasm has remained unclear. In this study, we showed that sensing of *in vitro* HCMV infection in primary fibroblast cells is largely driven by recognition of exogenous DNA, rather than direct sensing of the viral genome.

Our findings have important implications for studies of innate immune sensing using *in vitro* infection systems. Many reports investigating sensing of HCMV rely on standard laboratory virus preparations similar to those used here. Our results demonstrate that exogenous DNA contamination can act as a dominant confounding factor in fibroblast infection models. Careful consideration of virus preparation methods, including nuclease treatment and rigorous controls, is therefore essential to accurately dissect virus sensing mechanisms. Importantly, our data does not exclude cGAS as a sensor of HCMV. Indeed, a recent study demonstrated inhibition of cGAS within the nucleus by viral lncRNA4.9, suggesting nuclear sensing of HCMV genomic DNA [23]. Additionally, low levels of cytoplasmic detection (due to capsid leakage) may still occur in absence of exogenous DNA; however, this is likely inhibited by viral antagonists of cGAS, such as pp65 [24, 25].

More broadly, these results highlight a general caveat for studies of DNA virus sensing *in vitro*, where host or extracellular DNA, co-purified during standard virus production, can artificially amplify cGAS-dependent responses. Similar issues may extend to other viruses and experimental systems, contributing to discrepancies between *in vitro* and *in vivo* observations of innate immune activation.

## Materials and methods

### Cell culture and virus

Human dermal fibroblast (HDFn) cells obtained from Thermo Fisher Scientific were cultured in DMEM (Sigma-Aldrich, D5796) supplemented with 10 % FBS at 37 °C with 5 % CO_2_. HEK-Blue IFN-α/β cells obtained from InvivoGen were cultured in DMEM supplemented with 10 % FBS, 30 µg/ml blasticidin and 100 µg/ml zeocin.

### RNA interference

cGAS-targeting siRNA was obtained from Integrated DNA Technologies as single-stranded oligonucleotides (sense: GGAAGGAAAUGGUUUCCAAdTdT; antisense: UUGGAAACCAUUUCCUUCCdTdT)[12] and annealed in-house. Subconfluent HDFn cells seeded on 12-well formats were transfected with 40 pmol/well of siRNA using Lipofectamine RNAiMAX (Invitrogen, 13778-150) per manufacturer’s instructions 48 h before infection.

### Virus stock preparation

BACs for TB40/E-GFP and AD169 were gifts from Felicia Goodrum [26] and Thomas Shenk [27], respectively. Master stocks were produced by nucleofection of 10^6^ HDFn cells with 4 µg BAC using program “T-016” on Amaxa Nucleofector II according to manufacturer’s protocol. Cells were expanded to T175 flasks following plaque formation. After 14 days, supernatant and cells were harvested and subjected to one freeze-thaw cycle. Virus was propagated in HDFn cells at an MOI of 0.1 until near 100 % cytopathic effects (12-16 days). Virus was harvested and propagated as before to produce the working stock. Supernatant and cells were harvested and cell-associated virus released using a Dounce homogeniser. Debris was removed by centrifugation at 1500 x g for 20 minutes at 4 °C. Cell-associated virus and supernatants were concentrated through 20 % sorbitol cushions via centrifugation at 24,000 rpm for 2 h at 4 °C (Beckman Coulter Optima L100-K, rotor: SW32Ti). Virus was resuspended in DMEM supplemented with 10 % FBS. The process was repeated to generate mock virus from uninfected cells.

### DNase treatment of virus stocks

Virus stocks were treated with 0.08 U/µl of TURBO DNase (Invitrogen, AM2238) for 90 minutes and enzyme removed using inactivation beads per manufacturer’s protocol. After treatment, subsamples were used for DNA quantification by Qubit dsDNA BR assay kit (Invitrogen, Q33266) per manufacturer’s protocol, for plaque assays, and in infection experiments.

### Plaque assay

Confluent HDFn cells seeded on 24-well plates were inoculated with serially diluted virus and incubated for two hours with occasional agitation. After incubation, cells were overlayed with immobilisation media (DMEM supplemented with 0.5 % Carboxymethyl cellulose and 2 % FBS).

### Infections

Virus was diluted in serum-free media according to indicated MOIs and used to inoculate confluent HDFn cells seeded on 12-well plates. Two hours post-infection, inoculum was replaced with DMEM supplemented with 10 % FBS.

### RT-qPCR

RNA was extracted using TRIzol reagent (Invitrogen, 15596026) and genomic DNA removed using TURBO DNase (Invitrogen, AM2238), both per manufacturer’s guidelines. cDNA was produced using High-Capacity cDNA Reverse Transcription Kit (Applied Biosystems, 4368814) and analysed by qPCR using TaqMan Gene Expression Master Mix (Applied Biosystems, 4369016). Primer-probe mixes were procured from Applied Biosystems (cGAS: Hs00403553_m1, IFNB1: Hs01077958_s1, IFIT1: Hs01675197_m1, CXCL10: Hs00171042_m1, GAPDH: Hs02786624_g1). Data was normalised to GAPDH using the delta-delta Ct method.

### Cell-based assays for the detection of type I IFNs

Secreted type I IFNs were detected using HEK-Blue IFN-α/β cells per manufacturer’s instructions. Briefly, supernatants from IFN producing cells were transferred to 96-well tissue culture plates, combined with HEK-Blue IFN-α/β cells and incubated for 24 hours at standard tissue culture conditions before proceeding with SEAP detection using QUANTI-Blue solution (InvivoGen, rep-qbs). After incubation for 1 h at 37 °C, absorbance (OD = 620) was measured using the Cytation3 plate cytometer (BioTek).

### IF Microscopy

Cells seeded on coverslips were fixed with 4 % paraformaldehyde for 15 minutes at indicated times. Cell membranes were permeabilised using 0.01 % Triton X-100 in blocking buffer (3 % horse serum and 1 % bovine serum albumin in PBS). Cells were incubated overnight at 4 °C with antibodies against IRF3 (1:250, Cell Signaling Technology, 11904S) and HCMV IE1/IE2 (1:2000, CMV-IE, Millipore, MAB8131). For secondary staining, cells were incubated with anti-Rabbit Alexa Fluor 488 (1:1000, Invitrogen, A21206) and anti-Mouse Alexa Fluor 647 (1:1000, Invitrogen, A21235) for two hours at room temperature. DNA was stained with Hoechst 33342 (1:10,000, Thermo Fisher Scientific, H3570) for 10 minutes. Coverslips were mounted using Prolong Gold Antifade (Invitrogen, P36930). Images were captured using Zeiss AxioObserver Z1 and analysed in Zeiss Zen Lite.

## Acknowledgements

This work was funded by the Biotechnology and Biological Sciences Research Council BBS/E/RL/230002A to FG.

